# The effect of tongue-tie application on stress responses in resting horses

**DOI:** 10.1101/634717

**Authors:** Laura Marsh, Paul McGreevy, Susan Hazel, Luiz Santos, Michelle Hebart, Samantha Franklin

## Abstract

Tongue-ties (TT) are commonly applied to both Standardbred and Thoroughbred racehorses to increase control, by preventing them from getting their tongue over the bit, and as a conservative treatment for equine respiratory conditions, principally dorsal displacement of the soft palate. This study investigated responses to TT application in horses, at rest, using both behavioural (head-tossing, ear position, gaping and lip licking) and physiological (salivary cortisol concentrations, eye surface temperature and heart rate) indices. Twelve Standardbred horses (six of which were naïve to TT) were used in a randomised crossover design. The study comprised 3 phases; Phase 1 (Baseline), Phase 2 (Treatment), and Phase 3 (Recovery). At phase 2, tongue tie application (TTA) was performed using a rubber band that was looped around the tongue and secured to the mandible for 20 minutes. The control treatment (C) incorporated 30 secs of tongue manipulation, at the start of the 20 min, however no TT was applied. Behaviours (head-tossing, ear position, mouth gaping and lip-licking) and heart rate (HR) were recorded for the duration of the study and analysed in ten minute intervals. Salivary samples were taken at the end of each phase for subsequent cortisol assays and infrared thermography images were taken of each eye at 5-minute intervals. Statistical analyses were performed in SPSS using linear mixed effects models and repeated measures general linear models, to determine differences between treatments and within treatments, over time. Compared to control, there was more head-tossing/shaking (p<0.001), gaping (p<0.001) and backwards ear position (p<0.001) and less forward ear position (p<0.001) during TTA, in Phase 2. Horses with previous experience of TT showed more head-tossing (p=0.040) and gaping (p=0.030) than naïve horses. Lip-licking was more frequent after TTA treatment than control, during Phase 3 (p<0.001). Salivary cortisol concentrations increased after TTA (1846.1pg/mL ± 478.3pg/mL vs 1253.6pg/mL ± 491.6pg/mL, p=0.047). Mean HR, and mean right and left eye temperature did not differ significantly between treatments in any phase (all p> 0.05). The findings of this study suggest the application of a tongue-tie causes changes to both behavioural and physiological parameters suggestive of a stress-related response. Further research is needed that will enable racing and sport horse regulatory bodies to make informed decisions about the appropriate use of tongue-ties in horses.

## Introduction

The tongue-tie (TT is a form of tack modification that has been used in horses for over 100 years (Fleming, 1889). The device is used to hold the tongue in a fixed position and may be made from a rubber band, leather strap, nylon stocking or similar material, that is tied below the jaw or at either side of the horse’s mouth to the bit, after being looped around the tongue. Early reports suggest that TTs were used to prevent abnormal noise and airway obstruction arising as a result of the horse “retracting the tongue so as to force back the soft palate to such an extent that it interferes with the passage of air between the nasal passages and the larynx” (Fleming, 1898). Today, TT use remains commonplace in both Standardbred and Thoroughbred racehorses, throughout the world (1–6). Their primary putative purposes are firstly to conservatively address airway patency issues, principally due to dorsal displacement of the soft palate (DDSP) and improve performance and secondly, to aid control of the horse by preventing it from getting its tongue over the bit (3, 6, 7). Dorsal displacement of the soft palate, one of the most common causes of dynamic airway obstruction during strenuous exercise is thought to affect approximately 20% of racehorses (Pollock et al. 2009; Priest et al., 2012.). Tongue-ties are frequently used as the first line of conservative treatment by trainers and may also be used in combination with surgical management (Franklin et al., 2001; Barakzai and Dixon, 2005; Barakzai et al., 2009a). However, the efficacy and the exact mechanism by which the TT may aid in prevention of DDSP remains controversial (Beard *et al.* 2001; Cornelisse *et al.* 2001; Franklin *et al.* 2002;(5, 8).

Over recent years, concerns regarding potential welfare issues associated with TT use have been raised by animal welfare organisations (Barakzai, 2009b). This has led to these devices being banned from a number of equestrian disciplines under *Federation Equestre Internationale* regulations (9). Anecdotal reports suggest that routine TT application may cause damage to the tongue (including lacerations, dysphagia, bruising, swelling, discolouration and paralysis) (10). A recent South Australian survey identified that 26.3% of Standardbred racehorse trainers reported complications relating to tongue-tie use, mostly associated with swelling of and superficial cuts to the tongue, as well as changes in behaviour including head shyness (11). A study in the UK also reported that tongue-ties were not well tolerated in young Thoroughbred racehorses (12). This implies that the horses must habituate to the aversiveness of the procedure (McGreevy and McLean, 2010). It is not clear how long horses take to habituate to TTs and whether they ever do so completely. Whenever sustained pressure is used to modify horse behaviour, the principles of ethical training as espoused by the International Society for Equitation Science (ISES 2011) are compromised and negative reinforcement that relies on the *release* of pressure, cannot take place. The application of various training devices in horses, including bit attachments and restrictive nosebands, have been reported to result in pain and stress responses, thus compromising welfare (13–15).

Appropriate assessment of stress in animals involves the integration of measurement of both behaviour and physiology. Changes in behaviour can provide a useful and immediate means of assessing the response of an animal to its environment, and shifts in demeanour, posture, gait and interactive behaviour may be associated with presence of pain or stress (16). Head behaviours (such as head-shaking/-tossing and ear positioning) as well as oral behaviours (gaping and lip-licking), have been used to estimate an animal’s affective state (17). To date, the physiological assessment of stress in horses has been based primarily on changes in endocrine function (18), as well as parameters that reflect changes in autonomic functioning including heart rate and eye temperature (ET) (18–20). Salivary cortisol concentration is established as a non-invasive indicator of stress because it reflects activation of the hypothalamo-pituitary-adrenocortical axis (HPA) (21, 22). Measurement of maximum ET using infrared thermography has the potential to assess both acute and chronic stress in animals, reflecting changes to peripheral blood flow during increased sympathetic output (22–25). Heart rate variability (HRV) is a measure of autonomic function and can be used to determine the balance between sympathetic and parasympathetic tone (20, 24, 26, 27).

The effect of tongue-tie application on stress responses in horses has not yet been investigated. The aim of this study was to determine the effect of TT application on stress responses in resting horses. It was hypothesised that the application of a TT would induce a stress response, resulting in increased concentration of salivary cortisol, ET, HR and conflict/agitated behaviours. We also hypothesised that the stress response would be exacerbated in horses that were naive to TT application.

## Materials and Methods

### Animals and husbandry

This study used Standardbred horses (n=12), comprising mares (n=8) and geldings (n=4) aged 11.5 ± 3.0 years (mean ± s.d) and weighing 487 ± 33.9 kg (mean ± s.d). Animals were subjected to a health check prior to the trial, and deemed free from illness or injury. On experimental days, horses were brought in from the paddock, and housed in day yards before being secured in stocks during the experiment. Horses wore a halter only, with no bit and were loosely tied with a rope to the side bar of the stocks. All procedures were approved by the University of Adelaide Animal Ethics Committee (S-2015-141, 02/07/2015), and performed in accordance with the *Australian Code of Practice for the Care and Use of Animals for Scientific Purposes*, 8^th^ addition (2013) (28).

### Experimental Design

The study was based on a randomised crossover design comprising two treatment groups: tongue-tie application (TTA), and control (C). Horses were classified as either naive (n=6) or having previous experience of TTA during their racing careers (n=6). Horses were pseudo-randomised and assigned to treatments so that 6 horses (3 experienced and 3 naïve to TT), would have TTA on day 1 followed by C on Day 2, and 6 horses (3 experienced and 3 naive), received C on Day 1 followed by TTA on Day 2. For each horse, treatments were performed at the same time on both days to take account of diurnal rhythm in physiological parameters. For each treatment, horses were observed for a total of 80 minutes, which was divided into three phases (Phase 1: 30-minute baseline; Phase 2: 20-minute treatment; and Phase 3: 30-minute recovery). The application of a TT was performed by the same operator, using a rubber band measuring 152mm × 15mm (Belgrave rubber band; size 106; www.quillstationary.com.au). Following industry practice, the band was looped twice around the tongue, and then secured to the lower jaw by looping the TT around the mandible (Figure 1). A new tongue-tie was used for each horse. The duration of 20 minutes was based on the median duration of TT use reported during training and racing in a previous study (Findley *et al.*,2015). During C, tongue manipulation was performed for 30 seconds at the start of phase 2. This involved grasping of the tongue to mimic the initial process required for application of a TT, however no TT was placed. On each experimental day, horses were brought into the barn in pairs, and individually restrained within stocks for 10 minutes to acclimatise to the surroundings and to allow instrumentation of the horses. During each treatment period, as one horse was treated, the other stood by, as a companion.

**Figure 1.**
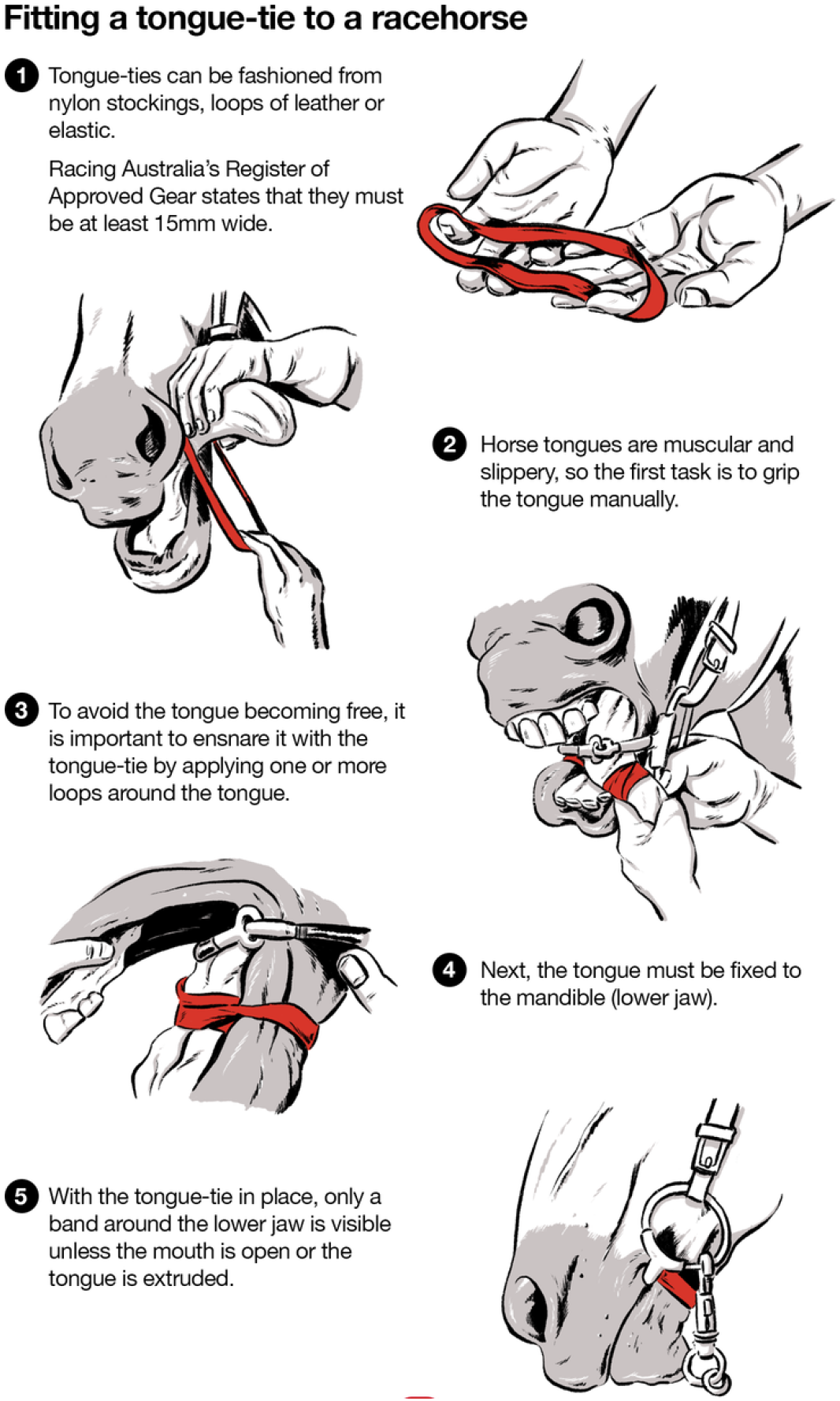
Horse with elastic tongue-tie in place. In alignment with industry practice, the device is looped twice around the tongue, and then the tongue secured to the lower mandible. NB: The horses in the current study did not wear bits. (Illustration from (29), republished under Creative Commons licence.)

### Behavioural Measures

Behavioural data were recorded using two digital cameras (GoPro® HERO3), each mounted on a tripod placed at either side of the stock at 90° to the horse’s head, at a distance of 1.5m. Recording commenced at the start of Phase 1 and ceased after completion of Phase 3. The behaviours recorded from the video record included oral (gaping, lip-licking) and head-related behaviours (head-tossing, ear positioning), that have previously categorised as either positive/relaxed or negative/agitated (Table 1) (17). Behaviours were measured as a duration (% of time), apart from lip-licking which was measured as a frequency (n). Video clips of ten minute duration (last ten minutes of phase 1 all of phase 2 and the first and last 10 minutes of phase 3) were used for behavioural analysis using behaviour analysis software (Mangold International GMbH, version interact 8) (Table 2). Analysis could not be blinded because it was not possible to obscure the observer’s view of the tongue. All behavioural analysis was conducted by the same observer to minimise inter-observer variation.

**Table 1:** Head-related and oral behaviours recorded in horses during and after tongue-tie application or tongue manipulation. Behaviours were categorised as either positive/relaxed or negative/agitated (17).

| Behaviour | Category | Description | Measurement |
| --- | --- | --- | --- |
| Head-tossing | Agitated/<br>conflict | Repeated throwing/tossing of the head either in an dorsal, ventral or lateral direction | Duration (%) |
| Ears back | Agitated | Ears were in a backwards position; both pinnae facing backwards | Duration (%) |
| Ears forward | Relaxed | Ears were in a forward position; both pinnae facing forward | Duration (%) |
| Lip-licking | Ambiguous | Protrusion of the tongue to lick the lower or upper lips. | Frequency (n) |
| Gaping | Conflict | Repeated opening of the mouth such that space is visible between upper and lower jaw | Duration (%) |

**Table 2:** Descriptions of the phases and time points for analysis of behaviours.

| Phase | Time point (minutes) | Description |
| --- | --- | --- |
| 1 | 20-30 | Baseline |
| 2 | 0-10 | T1 |
| 2 | 10-20 | T2 |
| 3 | 0-10 | Post1 |
| 3 | 20-30 | Post2 |

### Physiological measures

#### Salivary Cortisol

Saliva samples were collected using Salivette® (Sarstedt, Sweden) swabs, at the end of each phase. The swab was placed in the horse’s oral cavity (between the cheek and the teeth) using surgical forceps and moved around gently for 30 seconds. Swabs were stored on ice before being transported to the laboratory, where the tubes were centrifuged (1000g for 10 minutes) and samples frozen at −20°C until later analysis. The saliva was analysed for cortisol concentration, using a commercially available ABOR Enzyme-linked immunosorbent assay (ELISA) (Ann Arbor, Michigan 48108-3284 United States). Sensitivity and limit of detection were determined to be 17.3 pg/mL and 45.4pg/mL, respectively. All samples were run in duplicate and results expressed as pg/mL. The intra-assay coefficient of variation determined from duplicates of a control saliva sample in each assay plate (n=3) was 16.7% and the inter-assay coefficient of variation was 5.9%.

#### Eye temperature

An infrared camera (ThermaCam S60, FLIR Systems AB, Danderyd, Sweden) was used to collect thermographic images of the eye at 5-minute intervals for the duration of the study period, as per previous studies (Yarnell et al., 2013). At each time point, three images were taken of the right and left eye, and the clearest of the three used for analysis. Images were taken 90° from the front of the horse at a distance of 0.5-1.0 meters. Atmospheric temperature and relative humidity were recorded every 30 minutes from the Bureau of Meteorology (BOM) Roseworthy weather station, and values calibrated during image analysis. Maximum eye temperature (^0^C) was determined using FLIR Tools version 5.6, (FLIR Systems AB, Danderyd, Sweden), with the analysis of temperature taken from within the medial posterior palpebral border of the lower eyelid and lacrimal caruncle (Figure 2).

**Figure 2:**
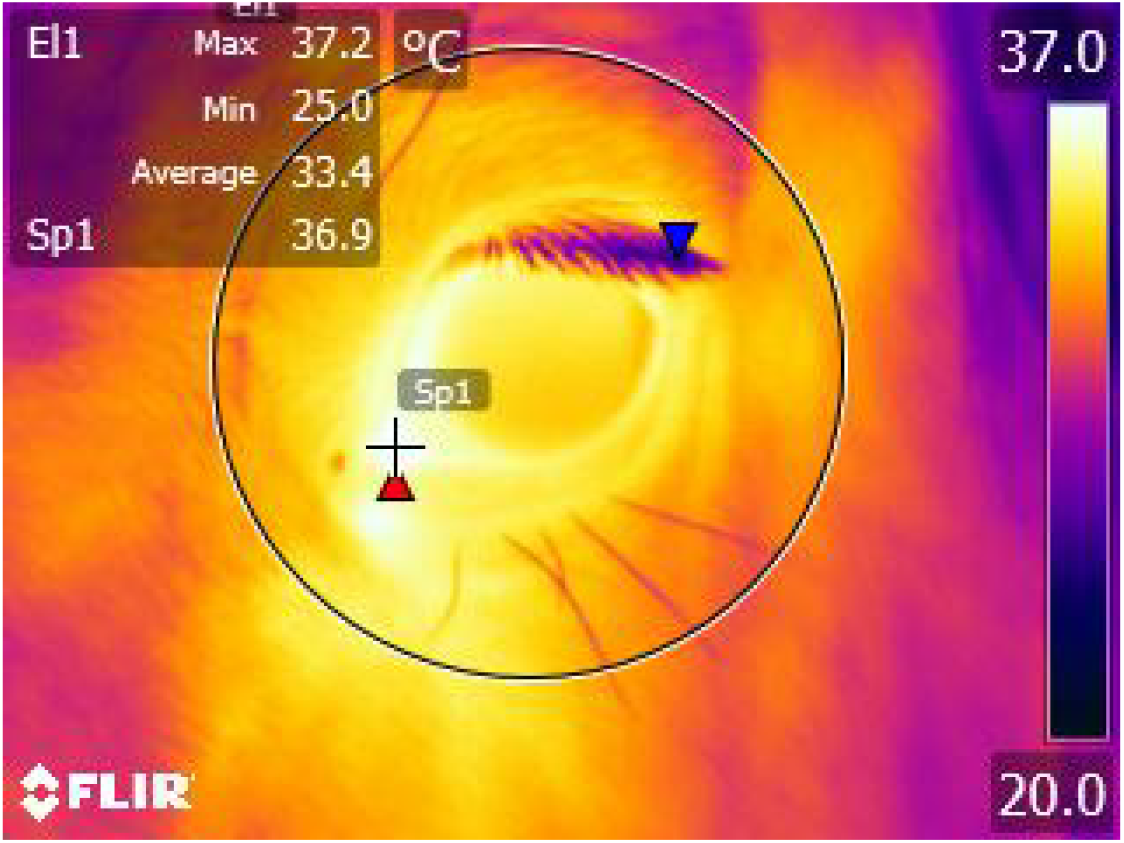
Thermographic image taken of a horse’s left eye. Maximum eye temperature was taken from within the medial posterior palpebral border of the lower eyelid and lacrimal caruncle.

#### Heart Rate

Heart rate was recorded using a telemetric ECG system (Televet 100; Engel Engineering Service GmbH; 63150 Heusenstamm, Germany). Adhesive electrodes were attached to the skin of the thorax in a modified base-apex configuration (with the left arm (+) and left leg (+) electrodes placed on the thorax, over the apex of the heart and the right arm (-) and reference electrodes placed over the left shoulder). These were connected to the Televet unit, which was fixed to a surcingle. ECG data was recorded onto a SD card and uploaded to a laptop computer after the conclusion of the test. The Televet software was used to calculate RR intervals (using a cut off value of 20% maximum deviation of consecutive RR intervals). ECG traces were checked and artefacts corrected manually prior to subsequent analysis.

#### Statistical Analysis

Statistical analysis was performed using SPSS 23 (SPSS Inc., Chicago, IL, USA) Data are reported as mean ± SEM. Data were tested for normality and homogeneity using the Kolomogorov-Smirnov and Levene test, respectively. Non-normally distributed data were log10 transformed. A linear mixed-effects model was used to analyse mean behaviour, salivary cortisol concentration, eye temperature and heart rate. Behaviour data were analysed from the last 10 minutes during phase 1, all of phase 2 and the first and last 10 minutes of phase 3. The model included fixed effects (treatment type, day, time, previous experience, sex and treatment sequence), and random effects (Horse ID). Baseline measurements for mean salivary cortisol concentration, eye temperature and heart rate were included as a covariate due to high inter-horse variability at baseline. A repeated measures general linear model was used to determine differences within treatments over time for both behaviour and physiological parameters, using within-subject effects of time, and between-subject effects as treatment. Variables were considered significantly different when p <0.05.

## Results

### Behavioural assessment

#### Head-tossing

Horses spent significantly more time head-tossing during TTA than C in both T1 (21.3 % ± 3.6 % vs 1.1 % ± 0.4 %; p = 0.001), and T2 (26.0 % ± 3.7 % vs 0.6 % ± 0.2 %; p = 0.001) (Figure 3). A significant association was observed between previous TT experience and head-tossing behaviour, in that horses with previous exposure to TT tossed their heads for longer (26.7 % ± 5.6 %) at T1 than naive horses (16.0 % ± 3.6 %) (p=0.04; Figure 4). No other factors had a significant effect on head-tossing.

**Figure 3:**
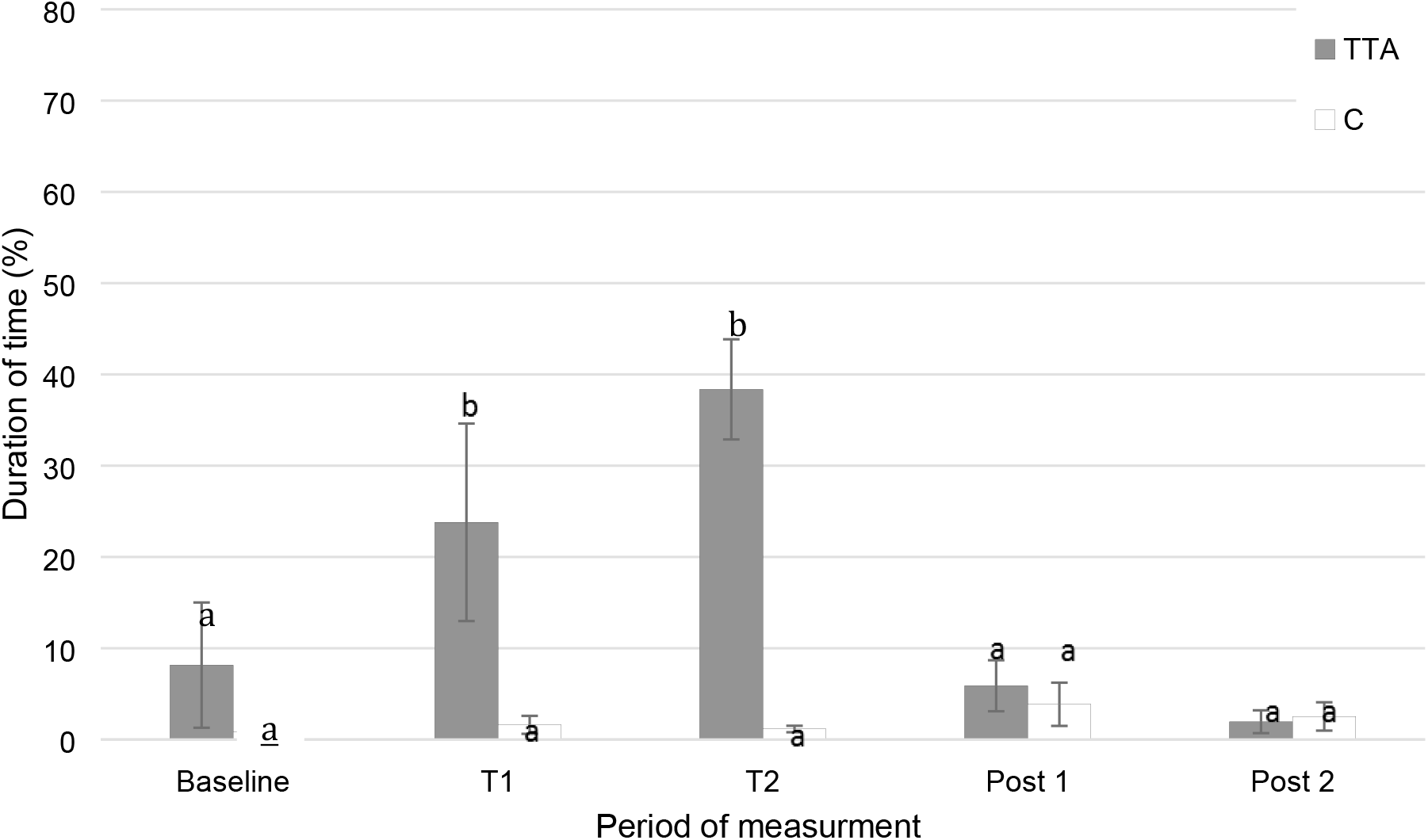
Mean (±SEM) percentage of time spent displaying head-tossing behaviour (%) for TTA and C groups over time. Values indicated with different subscripts (a, b) are significantly different (p<0.05). Key: Baseline = 10 minutes before treatment; T1 = first 10 minutes of treatment phase; T2 = last 10 minutes of treatment phase; post 1 = first 10 minutes of recovery; post 2 = last 10 minutes of recovery)

**Figure 4:**
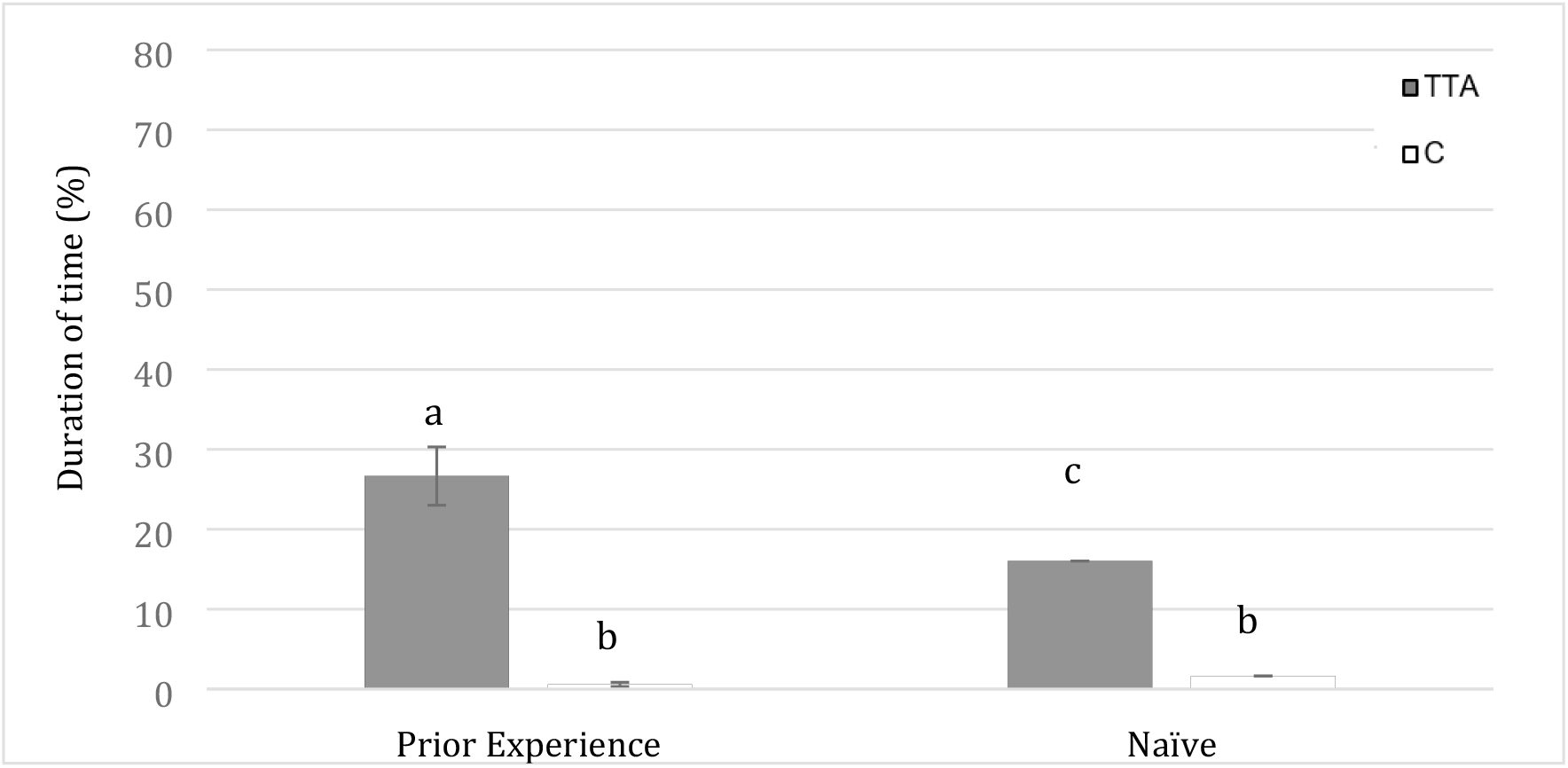
Mean ± SEM time spent displaying head-tossing behaviour (%) of naïve vs horses with prior experience of TT during TTA and C. Values with different subscripts (a,b,c) are significantly different (p<0.05).

#### Ear position

Horses undergoing TTA spent significantly more time with their ears in a backwards position than C at three time points: T1 (62.6 % ± 4.8 % vs 9.4 % ± 2.1 %; p<0.001), T2 (74.7 % ± 4.6 % vs 9.3 % ± 2.7 %; p<0.001) and Post 1 (18.9 % ± 6.6 % vs 9.0 % ± 2.1 %; p= 0.021) (Figure 5). Correspondingly, there was a significant difference in the time spent with their ears forwards in C compared to TTA at T1 (27.5 % ± 3.3 % vs 3.1 % ± 0.8 %; p<0.001) and T2 (29.3 % ± 5.2 vs 3.4 % ± 0.7 %; p<0.001).

**Figure 5:**
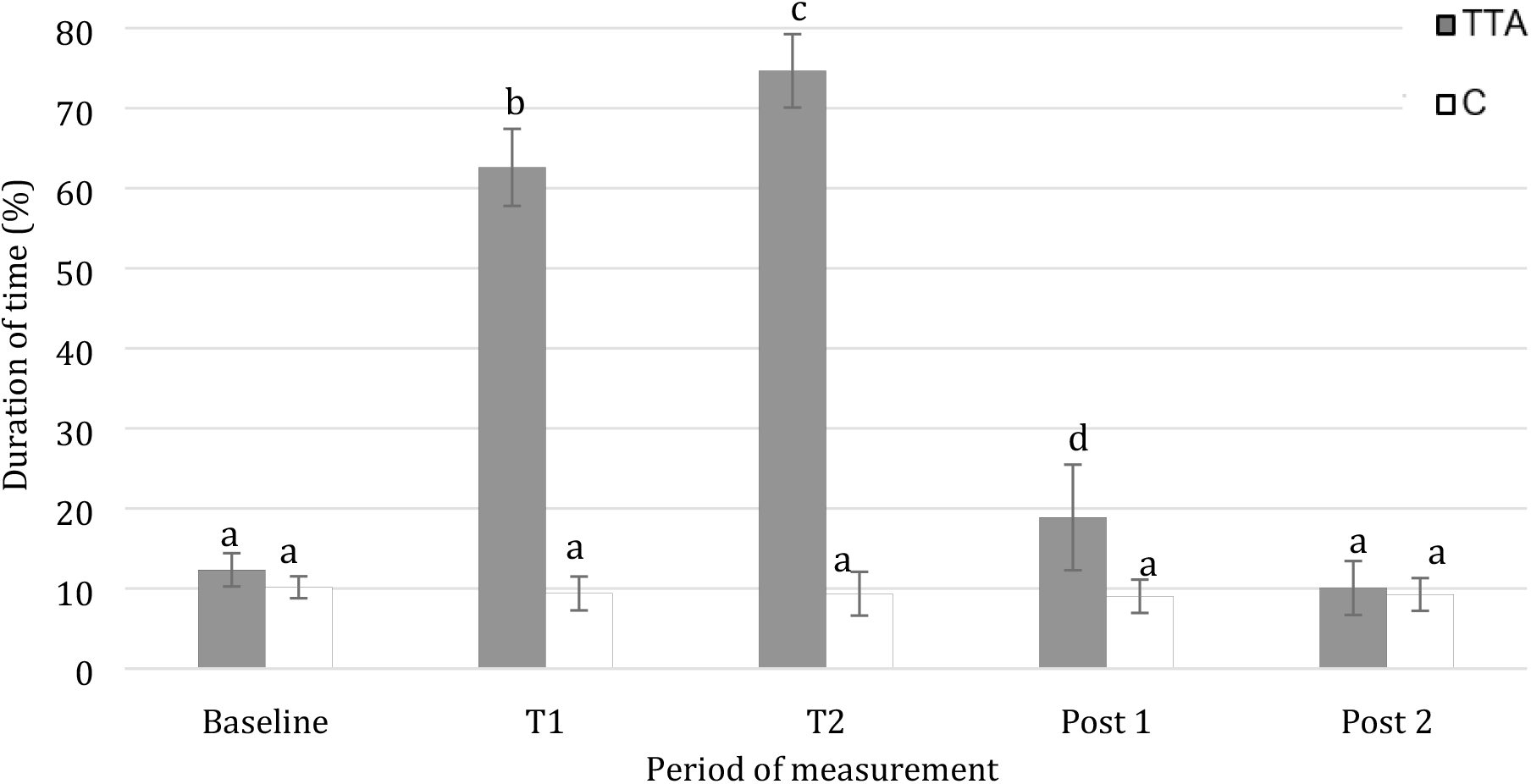
Mean duration of ears positioned backwards of horses (n=12) in TT and C throughout trial. ^a-b-c-d^ Values indicated with different subscripts are significantly different (p<0.05).

Horses spent more time with their ears backwards with increasing time wearing the TT: there was a significant difference between T1 and T2, (p<0.001) (figure 5). Once the TT was removed the time spent with ears in a backwards position was significantly decreased compared with T1 and T2 (P <0.001) (Figure 5).

Time of testing also had an effect at T2, with afternoon-tested horses spending more time with their ears backwards than morning-tested horses (47.8 % ± 7.1 % vs 37.8 % ± 6.5 %; p= 0.041). At Post 1, horses spent significantly more time with ears backwards on Day 1 compared to Day 2 (19.0 % ± 6.6 % vs 9.2 % ± 2.2 %; p= 0.025). Accordingly, during Post 1, horses spent less time with ears facing forward on Day 1 compared to Day 2 (24.4 % ± 3.3 % vs 32.1 % ± 2.4 %; p= 0.010).

An interaction between previous experience and treatment showed horses with prior experience of tongue-ties spent less time with ears forward compared to naive horses at Post 1 (21.3 % ± 6.5 % vs 28.5 % ± 3.3 %; p=0.032). No other factors had a significant relationship with ear position

#### Gaping

No horses showed gaping behaviour during the baseline recordings. In C, no gaping was observed in any horse throughout the recording period. However, during TTA, horses spent significant amounts of time showing gaping behaviour both at T1 (58.2 % ± 3.4 % TTA vs 0.0 % ± 0.0 % C; p< 0.001) and T2 (48.9 % ± 3.7 % TTA vs 0.0 % ± 0.0 % C; p< 0.001) (Figure 6). Gaping declined significantly after TT removal (4.60% +/− 0.66%) p< 0.001 during post 1) and no gaping was observed by 20 minutes post removal.

**Figure 6:**
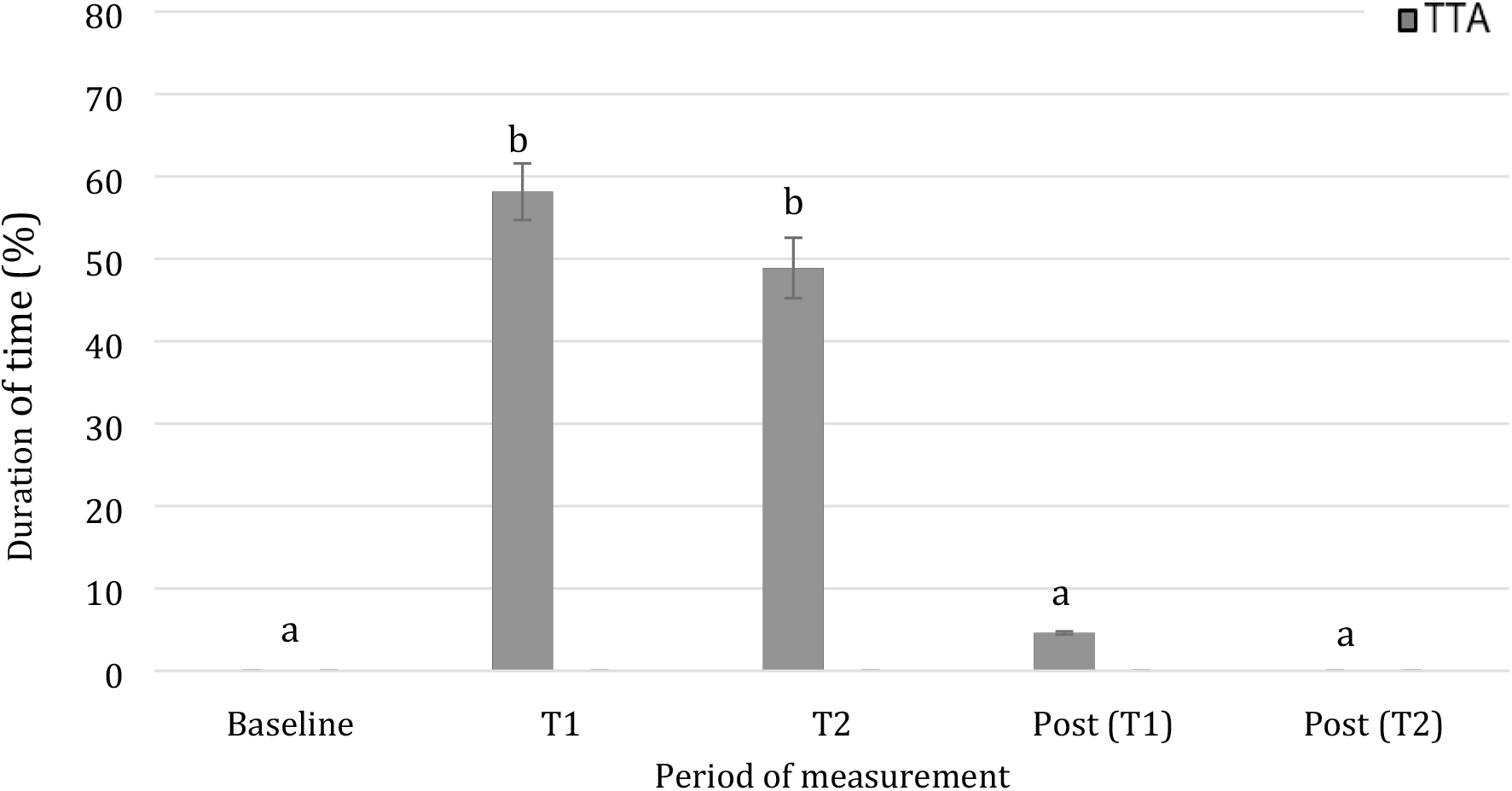
Mean duration of gaping behaviour of horses (n=12) during TTA throughout trial. ^a-b^ Values indicated with different subscripts are significantly different (p<0.05).

There was a significant interaction between previous TT experience and gaping behaviour: horses with previous experience of TT gaped more than naive horses at T2 (58.1 % ± 2.6 % vs 48.9 % ± 4.3 %; p= 0.030). No other factors had a significant relationship with gaping.

#### Lip-licking

Lip-licking was observed significantly more frequently in TTA than in C at Baseline (12.8 ± 2.7 vs 6.3 ± 1.7; p=0.014), Post 1 (32.9 ± 4.1 v 3.8 ± 1.40; p<0.001) and Post 2 (20.1 ± 2.2 vs 3.6 ± 1.0; p<0.001) (Figure 7). Lip licking was not possible during TTA due to the tongue being secured to the mandible. However, there was a significant increase in lip licking following TT removal at Post 1 (32.02 ± 14.29; p<0.001), and Post 2 (20.08 ± 7.63 TTA; p<0.001) compared to baseline (12.08 ± 2.31; p<0.001), Subsequently, there was a significant reduction in lip-licking between Post 1 and Post 2 (32.9 ± 4.1 vs 20.1 ± 2.2; p<0.001).

**Figure 7:**
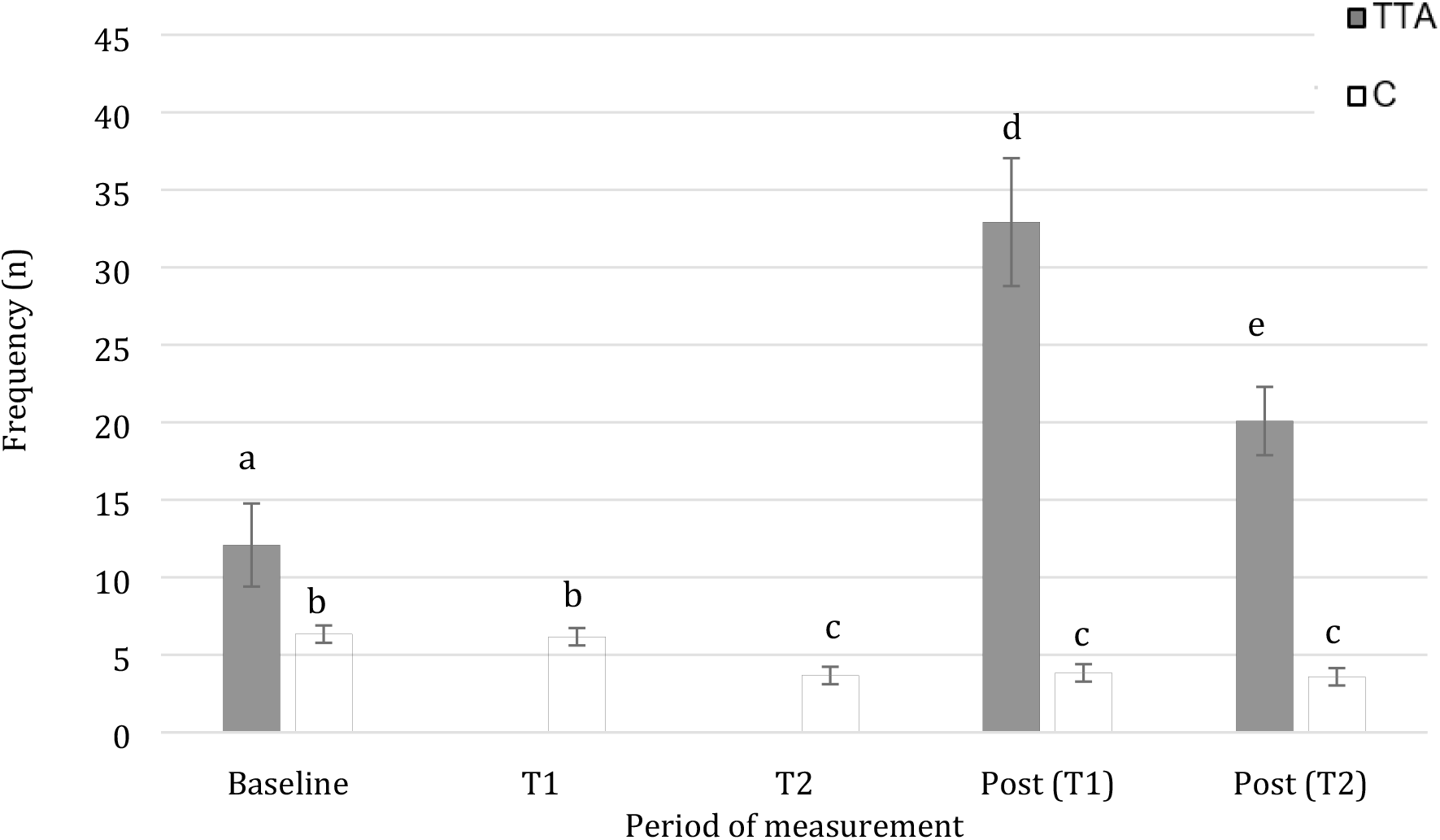
Mean frequency of lip-licking behaviour of horses (n=12) during TTA and C throughout trial. ^a-b-c-d-e^ Values indicated with different subscripts are significantly different (p<0.05).

### Physiological assessment

#### Cortisol

Salivary cortisol concentration did not differ between horses with TTA and C at Phase 1 (p= 1.29) or phase 2 (p= 0.89). A significant difference in saliva cortisol concentration was observed between horses at Phase 3, with horses showing increased salivary cortisol concentrations after TTA than compared to C (1846.1pg/mL ± 478.3pg/mL vs 1253.6pg/mL ± 491.6pg/mL; p=0.047) (Figure 8).

**Figure 8:**
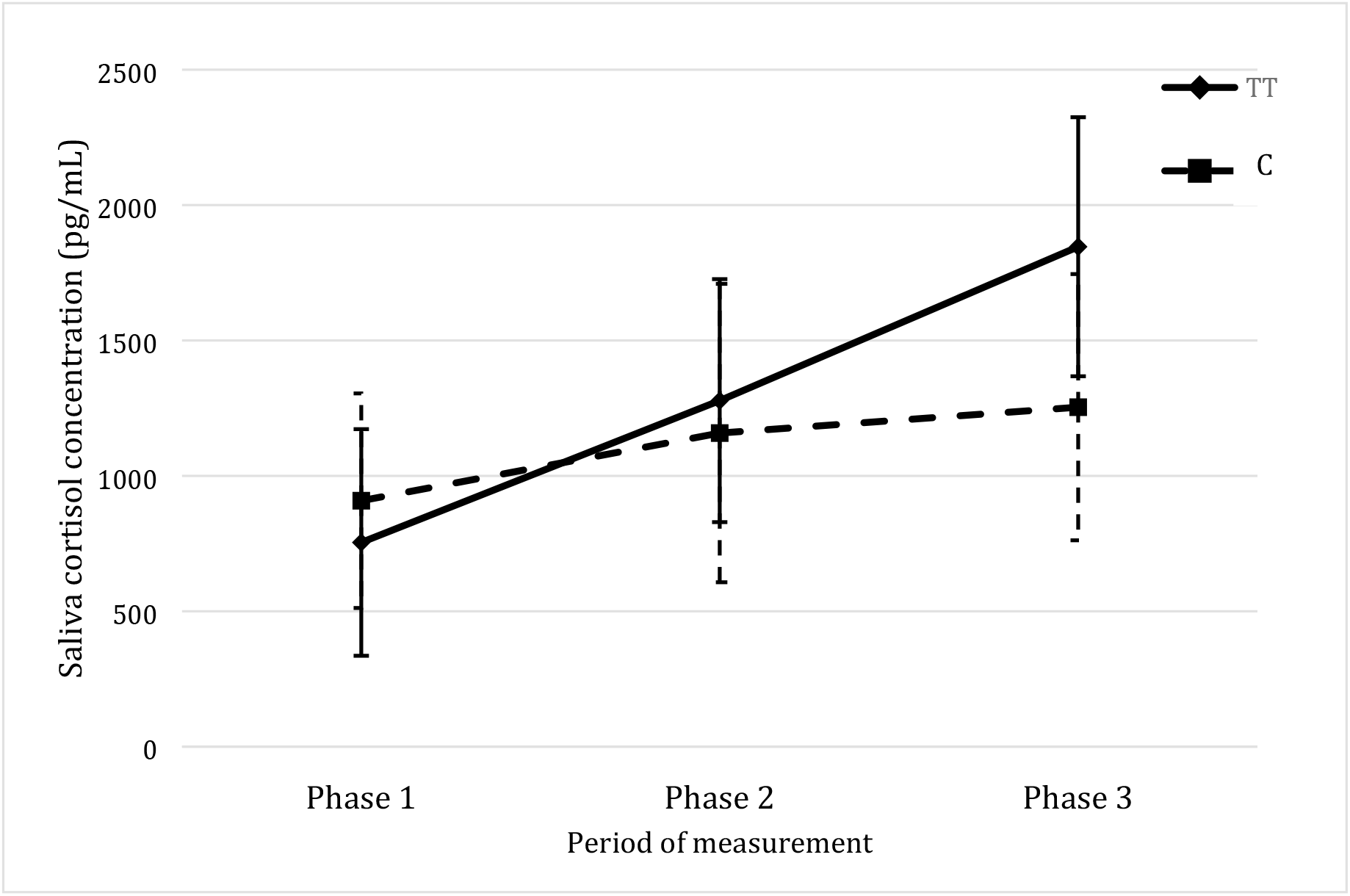
Mean saliva cortisol concentration (pg/mL) in TTA and C throughout trial.

#### Eye temperature

No significant difference in mean right and left eye temperature was observed between treatments during any phase of the experiment Phase 1 (36.1 ± 0.08°C TTA vs 36.2 ± 0.10°C; p=0.485 and 36.25 ± 0.10°C TTA vs 36.4 ± 0.13°C; p=0.463), Phase 2 (36.2 ± 0.16°C TTA vs 36.3 ± 0.10°C; p=0.190 and 36.3 ± 0.11°C TTA vs 36.5 ± 0.13°C; p=0.234) and Phase 3 (36.1 ± 0.08 °C TTA vs 36.4 ± 0.18°C; p=0.215 and 36.3 ± 0.09°C TTA vs 36.5 ± 0.07°C; p=0.369), respectively.

#### Heart rate

Mean HR did not differ significantly between TTA and C at any time point (Figure 9). However, there was a trend for horses in the TTA group to have a higher mean heart rate compared with C at T2 (43bpm ± 2.4bpm vs 37bpm ± 2.5bpm; p=0.079), followed by a reduction in HR in this group of horses after TT removal (figure 9)

**Figure 9:**
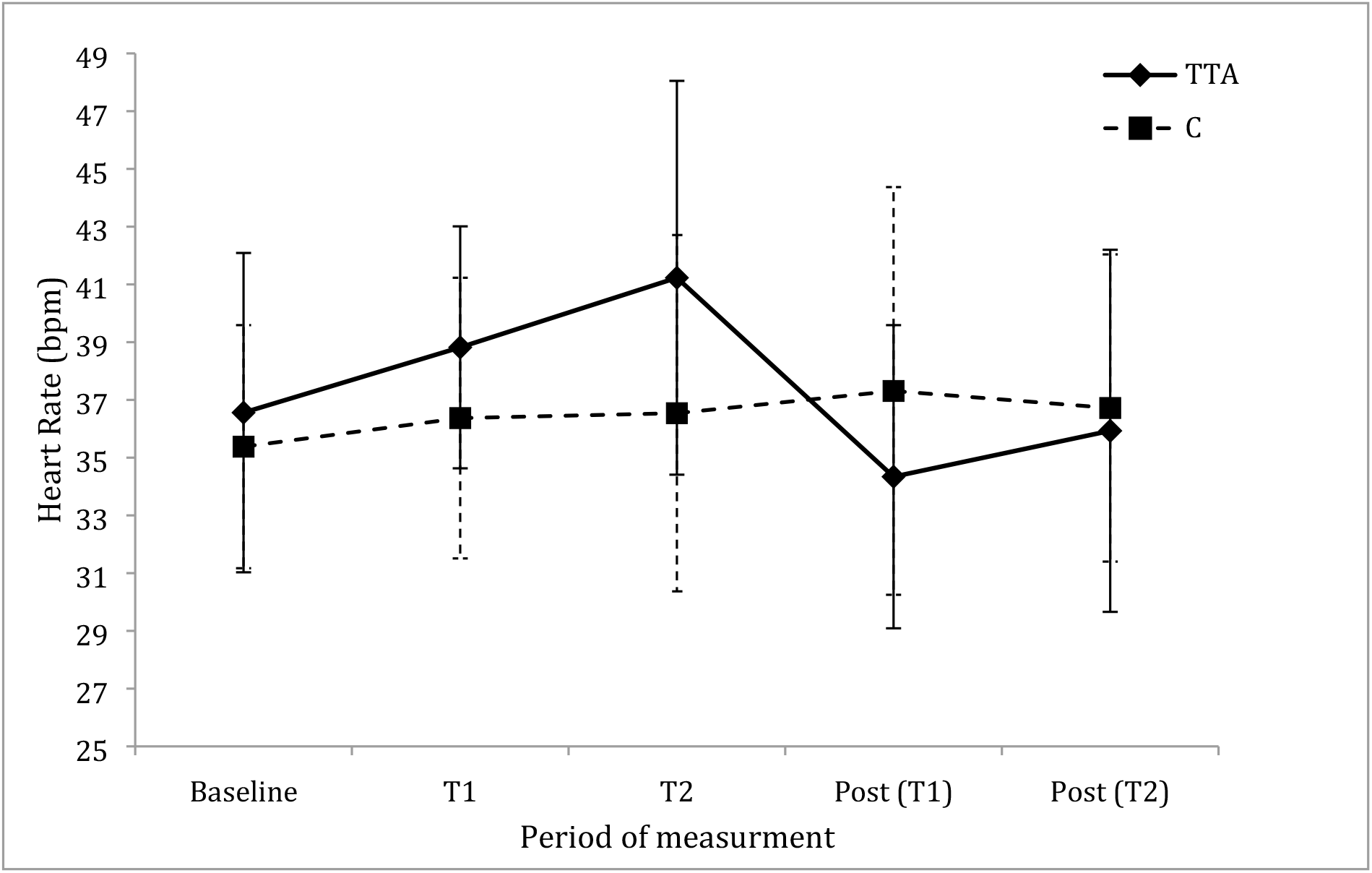
Mean heart rate (bpm) in TT and C throughout trial.

## Discussion

In this study, both behavioural and physiological indices were used to investigate possible indicators of stress in horses during and following TT application. The results support the hypothesis, suggesting TTA resulted in changes to behavioural and physiological parameters that may be indicative of a stress response.

### Behavioural responses

Behavioural responses to a discrete challenge can offer an immediate, easily identifiable and non-invasive means of assessing an animal’s affective state (17). Changes in the behaviours involving the head, ears and mouth have been reported to reflect horses in relaxed or agitated states (30). Conflict behaviour has been described as a response exhibited by animals that have difficulty coping with competing motivations associated with resistance to handling, training or equipment (31, 32). It is possible that horses wearing a TT are highly motivated to move their tongues and so they can be expected to attempt various oral and lingual manoeuvres to free themselves of the restraint.

### Head-tossing

Head-tossing behaviour in horses has been shown to reflect agitated or conflict behaviour caused by irritation or pain to the mouth (33). In the current study, horses exposed to TTA spent significantly more time head-tossing during Phase 2 compared to C, suggesting that the presence of a TT may be irritating and/or results in discomfort or pain, inducing a negative affective state in these animals. Horses with previous experience to TTA showed more head-tossing than those with no previous experience. A recent study found the application of TT was not well tolerated by naïve horses, confirming the need for habituation (12). Habituation of novel stimuli by means of repeated exposure to that stimulus, may allow animals to become accustomed to the environment and thus reduce fear responsiveness (34). However, in the current study, habituation did not take place during the treatments nor was there evidence of it having occurred before the treatments took place. Indeed, the frequency of head tossing behavior increased between T1 and T2 and prior experience of TT use resulted in more behavioral responses to them. Information on the horses’ most recent previous exposure to TT, including the duration or tightness of repeated exposure was not available for horses in this study. Therefore, future research should explore how horses’ attempts to remove TT decline over time. This is important because the eventual disappearance of attempts to remove TT may be seen as either habituation or mark the point at which horses reach learned helplessness (Hall et al 2007).

### Ear position

Backwards positioning of the ears is reported to reflect negative affective states, including fear, discomfort, submission, avoidance, pain or aggression in horses (35–37). In our study, horses spent more time with their ears backwards during TTA and immediately after removal compared to C. These findings align with those from a study of horses ridden with their necks in a hyper-flexed position, in that horses spent more time with ears backwards when hyperflexed than when they were not hyperflexed, possibly suggesting an association between hyperflexion of the neck and a negative experience (38). During Phase 2, horses with TTA spent more time with their ears backwards at T2 than at T1. This may suggest that over the course of the 20 minutes of the current treatment, the horses’ discomfort, submission, avoidance or discomfort increased. This is consistent with previous research showing that the intensity, duration and frequency of a stressor generally correlates with the intensity of the stress responses that manifest (39). During Post 1, the time spent with ears in a backwards position was also higher with TTA than C. This may suggest that horses subjected to TTA may still be recovering from the effects of treatment into what we anticipated to be a wash-out period.

When horses have their ears facing in forwards are often regarded being relaxed or in a positive affective state (40). Another study reported ears facing forward as a feature of horses displaying interest or pleasure towards experienced handlers in contrast to inexperienced handlers (41). During TTA in the current study, horses spent less time with their ears forward at T1 and T2. This may suggest that during TTA, horses are less relaxed than during C. As with head-tossing, time spent with ears forwards was associated with previous experience. Horses with prior experience of TTA showed decreased time with ears in a forward position compared to naïve horses. Holding the ears back is recognised as a sign of pain (42) so this finding could is suggest that they have learned to associate the sensation of having their tongues tied with a negative affective state. It may also suggest that the experienced horses have learned that, rather than being swiftly transient, the restraint persists for some time.

### Gaping

Gaping or opening of the mouth has been categorised as an agitated behaviour, where increases in this behaviour may reflect a negative affective state in horses (17). Gaping has been suggested be a reflection of horses showing escape or avoidance type behaviours in response to the presence of a bit (40, 43). In the present study, no bit was in place and horses did not perform gaping behaviour at any time point during C. Similarly, horses with TTA showed no gaping behaviour during Phase 1 or Post2. However, at Phase 2, there was increased gaping behaviour during TTA application. Some gaping behaviour was also observed in the 10 minutes after TT removal but this was significantly less than during TTA and was not significantly different to baseline or C. These findings suggest that when tongue-ties were in place, horses may have been performing gaping to avoid or escape the presence of tongue-tie. Similar, to other behavioural indices, gaping behaviour was influenced by previous experience to TTA, with, horses with previous experience to TT spending increased proportion of time performing gaping behaviour compared to naive horses.

### Lip-licking

The frequency of lip-licking behaviour during TTA showed a significant decline from Baseline. This probably reflects the fact that horses with tongue-ties are unable to physically perform lip-licking when their tongues are restrained. However, after TTA, horses showed significantly more lip-licking behaviour than at baseline which may represent post-inhibitory rebound behaviour. Post-inhibitory rebound behaviour is the term given to an increase in the expression of a behaviour following a period of restriction or deprivation (44, 45), that may indicate that the deprivation (of the opportunity to perform certain normal behaviours) has deleterious effects on welfare (44, 46). In a similar study that focussed on restrictive nosebands, a post-inhibitory rebound behaviour was observed for yawning, swallowing and lip-licking behaviour (23). That said, the authors of that study noted that post-inhibitory rebound behaviours were observed for all groups with varying noseband tightness, thus the horse’s response may have been related to the novelty of having had two bits in the mouth rather that noseband tightness alone. Importantly, in the current study, no significant difference in lip-licking behaviour emerged between horses experienced to TTA and the naive horses, suggesting the post-inhibitory rebound response observed was likely due to the application of tongue-tie rather than any novelty effect. No significant difference in lip-licking was observed in horses during C treatment between Phases 1, 2 and 3, suggesting that the brief tongue manipulation performed had no effect on the horses’ motivation to lick their lips.

### Physiological responses

When animals experience stressful stimuli, the body attempts to return to homeostasis by creating alterations in neural, endocrine, immune, hematologic and metabolic functioning (47). Non-invasive sampling procedures (including salivary cortisol, ET and HR), are considered more advantageous in some prey species, as the collection method requires minimal disturbance to the animal and reduces the likelihood of anticipatory or nonspecific stress responses (48, 49).

### Salivary cortisol

Elevations of the concentration of glucocorticoid hormone cortisol are regularly used as an indicator of stress (50). A positive correlation has been shown between salivary and plasma cortisol in horses (51). Salivary cortisol provides a non-invasive alternative to plasma cortisol analysis, as blood collection via jugular venepuncture or from jugular catheterisation can cause increase plasma cortisol concentration for up to 130 minutes (52). In the present study, there was no difference in salivary cortisol concentration in horses during C, suggesting that the saliva swabbing itself was not stressful for horses. Horses showed an increase in cortisol following TTA, suggesting that this training device posed enough discomfort to elicit a stress response. Although cortisol concentration did not differ between treatments at Phase 2, we would expect that the increase would be delayed as the time it takes free cortisol to increase in saliva post-stressor is approximately 30 minutes (53). Although baseline values were variable among individuals, the amount of change in cortisol after TTA is similar to previous studies that measured salivary cortisol in response to sham clipping in horses (25), as well as social isolation and the sound of fireworks (30). It is difficult to determine if salivary cortisol increased as a result from the inability to perform normal behaviours when TT is in place, from pain and stress, or a combination of both. However, the fact that TTA was enough to elicit a stress response, suggests that the training device may compromise welfare.

### Eye temperature

Changes in eye temperature occur as a consequence of increased body temperature, as well as changes to peripheral blood flow during increased sympathetic output (54). Images of the eye measured using an infrared thermographic camera have been reported to provide a non-invasive method of assessing stress in animals. Previous research in horses using infrared thermography has identified increased eye temperature in association with noseband tightness (23), exercise (22, 24) and fear tests (55). However, others have reported that eye temperature is a poor estimate of core body temperature and is not correlated with accepted measures of stress including heart rate and salivary cortisol concentrations (Soroko et al., 2016). The results of this study found no significant difference in eye temperature between treatments, over time. Factors that may affect the degree of heat emitted from a surface include sunlight, distance, temperature, humidity, and wind (54). All of these factors were included in the image analysis however horses were loosely tied in stalls to allow for freedom for horses perform oral and head behaviours. Given that all horses performed some degree of head-tossing during the trial, it is possible that when taking images of the eye, the required distance of 0.5-1meter was not always met and this may have influenced the results.

### Heart rate

Changes in heart rate have previously been used as a non-invasive measure stress in horses (20, 24, 26, 27). Heart rate is regulated by both the sympathetic and parasympathetic branches of the autonomic nervous system (ANS). Increased heart rate, due to an increase in sympathetic activity, has been associated with a number of husbandry practices including branding procedures, restraint, transport, and social isolation (50, 56) In this study, no significant difference was observed in HR between TTA and C groups over time. However, there was a trend for heart rate to increase atT2 with TTA and to decrease following TT removal. The lack of significant findings may relate to the low number of horses used in this study and the high variation in HR between individuals. In addition, fluctuations in HR are labile and subject to both external and psychological influences (57) and may have been affected by external environmental factors (extreme weather conditions, construction sounds and traffic noise) that occurred on some days but not others. Other studies have used heart rate variability (HRV) as a non-invasive measure of stress in horses (58, 59)). However, this was not deemed appropriate in this study due to the high prevalence of 2^nd^ degree AV block that was observed. 2^nd^ degree AV block has been shown to significantly influence HRV variables when based on RR intervals (60).

### Limitations and future research

Although behaviour offers an immediate means of assessing an animal’s response to potential stressors, the interpretation of behaviours is often subjective between assessors. Therefore, future studies may benefit from assessing the inter- and intra-observer reliability of the behaviours measured in this study, and thus reducing limitations due to observer bias. This is particularly important in prey species (including the horse), as outward behavioural signs of fear and distresses may be masked as a means of survival (25). The cortisol assay produced large values for both inter and intra coefficient of variation, consequently limiting the reliability of results. This may have been due to error associated with pipetting or contamination from feed material within the samples. Heart rate measurements showed large variation between individuals in this study and may have been influenced by extraneous factors in some horses. This study was also limited by the small sample size. A larger sample size would reduce the variation among individuals and thus increase reliability of results.

## Conclusion

This present study provides novel evidence on the effects of tongue-tie application on stress responses in resting horses, suggesting the application of a TT results in increased agitated/conflict behaviour, decreased desirable relaxed behaviours, and increased salivary cortisol concentration. Further investigation into the appropriate use of TTs in horses should establish whether the costs to the horses, when wearing TT, are offset by the benefits to the horses and other stakeholders. This will enable racing and sport horse regulatory bodies to make informed decisions regarding the continued use of tongue-tie in horses.

## Acknowledgments

The authors would like to thank Kiro Petrovski for the use of the infrared thermographic camera and Sarah Weaver for her assistance with the ELISA assay.

